# Senolytic activity of small molecular polyphenols from olive restores chondrocyte redifferentiation and cartilage regeneration in osteoarthritis

**DOI:** 10.1101/686535

**Authors:** Marta Varela-Eirín, Adrián Varela-Vázquez, Carlos Luis Paíno, Antonio Casado-Díaz, Alfonso Calañas Continente, Virginia Mato, Eduardo Fonseca, Mustapha Kandouz, Alfonso Blanco, José Ramón Caeiro, María D. Mayán

## Abstract

Osteoarthritis (OA) is the most prevalent disorder of articulating joints and a leading cause of disability in humans, affecting half of the world’s population aged 65 years or older. Articular cartilage and synovial tissue from OA patients show an overactivity of the membrane channel protein connexin43 (Cx43) and accumulation of senescent cells associated with disrupted tissue regeneration. We have recently demonstrated the use of the Cx43 as an appropriate therapeutic target to halt OA progression by decreasing the accumulation of senescent cells and by triggering redifferentiation of osteoarthritic chondrocytes (OACs) into a more differentiated state, restoring the fully mature phenotype and cartilage regeneration. In this study we have found that small molecular polyphenols derived by olive extracts target Cx43 and senescence in OACs, synovial and bone cells from patients and in human mesenchymal stem cells (hMSCs). Our results indicate that these small molecules including oleuropein regulate the promoter activity of Cx43 gene. The downregulation of Cx43 expression by oleuropein reduce gap junction intercellular communication, cellular senescence in chondrocytes and enhance the propensity of hMSCs to differentiate into chondrocytes and bone cells, reducing adipogenesis. In concordance with these results, these small molecules reduce Cx43 and decrease Twist-1 activity leading to redifferentiation of OACs, which restores the synthesis of cartilage ECM components (Col2A1 and proteoglycans) and reduces inflammatory and catabolic factors IL-1β, IL-6, COX-2 and MMP-3 and cellular senescence orchestrated by p53/p21 together with the synthesis of SASP via NF-kB. Altogether, our results demonstrate the use of the olive-derived polyphenols such as oleuropein as potentially effective therapeutic agents to enhance the efficacy of hMSC therapy and to induce a pro-regenerative environment in OA patients by restoring cellular phenotype and clearing out senescent cells in joint tissues in order to stop or prevent the progression of the disease.

## Introduction

Osteoarthritis (OA) is the most common degenerative joint disorder characterised by progressive degeneration of synovial joints that lead to limited mobility and pain. OA is the major cause of disability with a major socio-economic impact, which affects an increasing number of the ageing population^1^. Cartilage damage is the most typical sign of OA. Yet, this condition also affects the whole joint including bone, synovium, muscles, tendons and ligaments^2^. Articular cartilage from patients with OA shows an accumulation of dedifferentiated and senescent cells^3–6^ together with an increase in the production of pro-inflammatory cytokines such as IL-1β and catabolic enzymes (matrix metalloproteinases (MMPs) and aggrecanases) that lead to the breakdown of cartilage extracellular matrix (ECM)^7^. The prevalence of OA is rising worldwide^8^ and the aetiology is still under study, with new insights into molecular mechanisms involved in OA progression beginning to open new possibilities to restore phenotypic stability of articular chondrocytes and to develop new approaches in order to promote cartilage repair and restore normal joint function in OA patients^9–12^.

Interestingly and consistent with other wound-healing disorders, osteoarthritic cartilage and synovial tissue from OA patients contain high levels of the gap junction protein connexin43 (Cx43). Therapeutic approaches targeting Cx43 are showing promise in benefiting several age-related and chronic degenerative diseases by modulating tissue regeneration, inflammation and response to injury^13,14^. Cx43 is linked to regeneration and response to injury through several different mechanisms, acting as a brake in wound healing in skin^14,15^, cartilage^4,16,17^ and neurodegenerative disorders such as Alzheimer’s disease^18^. Cx43 belongs to the integral membrane protein family called connexins which form hemichannels and gap junctions (GJs) that enable direct communication between neighbouring cells by interchanging electrical, metabolic and signalling molecules such as IP3, cAMP, ions, glutathione or siRNAs^19,20^. Additionally, these channel proteins act as scaffold proteins or signalling hubs via their cytoplasmic domains ^4,21^, regulating different key signalling pathways independently of their channel activity^4,22–24^. Several Cx proteins are expressed in developing and mature skeletal tissues ^19,25,26^. However, Cx43 is the major Cx protein expressed in chondrocytes, synovial cells and bone cells^10,12,27–29^ and it has been involved in normal development and function of joint tissues^25,30–32^, and in joint disorders including age-related bone loss^33^, rheumatoid arthritis^16,34^ and OA progression^10,12,25,27–29^.

During tissue regeneration and following injury, the dedifferentiation, redifferentiation and senescence processes play finely tuned temporal and spatial roles to reverse the loss of tissue in a precise way^35,36^. Cx43, via channel-dependent and -independent functions, has been involved in different phases of tissue regeneration including in controlling acute and chronic inflammation, cell differentiation, migration, proliferation or cellular reprogramming and lately in cellular senescence^22–24,37,38^. Results from our lab and others, demonstrated that Cx43 is a target of interest for the treatment of OA in order to stop cartilage degradation and to restore regeneration. Overexpression of Cx43 and enhancement of the GJIC in osteoarthritic chondrocytes (OACs) compromise the ability of these cells to re-differentiate, promoting a stem-like state by activating the chondrocyte-mesenchymal transition factor Twist-1 and p53/p21-mediated cellular senescence^4^. However we have recently demonstrated that this is a reversible loss, because downregulation of Cx43 reduces stemness, hypertrophy markers, senescence and therefore MMPs and proinflammatory mediators, and improves Col2A and proteoglycans’ levels in monolayer and 3D cultures^4^. Interestingly, this Cx43-sensitive circuit regulates joint inflammation, cellular plasticity and accumulation of senescent chondrocytes^4^. The downregulation of Cx43 in human osteoarthritic chondrocytes, using CRISPR technology and the GJ inhibitor carbenoxolone, restores chondrocyte redifferentiation and decreases the propensity of chondrocytes to undergo cellular senescence by downregulating Twist-1 and p53/p21/p16 and inhibiting the nuclear translocation of the master regulator of senescence secretory phenotype (SASP) NF-kB^4^. These results clearly indicate that Cx43 controls senescence and acts as a molecular switch in chondrocyte phenotypes within a wound healing process^4^. In fact, downregulation of Cx43 in different wound healing disorders halts disease progression by restoring tissue regeneration^39–43^.

In this study we describe the use of small molecules, based on olive phenolic compounds, to downregulate Cx43 in OA using 2D and human 3D cartilage models. We have found that oleuropein decreases Cx43 promoter activity and GJIC, thus enhancing osteogenesis and chondrogenesis in hMSCs and redifferentiation of OACs. Besides, downregulation of Cx43 by olive-derived small polyphenols in OACs reduces cellular senescence in chondrocytes, bone and synovial cells and the inflammatory and catabolic activity related with cartilage degradation in OA. Thus, these molecules by targeting Cx43 and senescence may be attractive candidates for tissue engineering and regenerative medicine strategies for OA treatment and for promotion of proper cartilage and joint repair in these patients.

## Results

### Olive-derived polyphenols including oleuropein impair adipogenesis and enhance the chondrogenic and osteogenic ability of hMSCs

Cx43, altered during OA^27^, is also a critical regulator of hMSCs differentiation^44–46^. Based on previous reported results from our group^4,27,30,47,48^, we used a small-scale screening to identify compounds that downregulate Cx43 (data not shown), and we have identified compounds such as the small molecule oleuropein (Fig. 1a). Oleuropein and an olive-extract (OE) containing 40% of oleuropein significantly reduced Cx43 protein levels in OACs treated with different concentrations for 2 hours, as detected by western-blot and by flow cytometry (Fig 1a). Importantly, polyphenols comprising oleuropein have been previously proposed and patented for the treatment of OA due to their ability to stimulate cartilage anabolism and prevent inflammation^49–51^. An MTT assay showed no effect of 0.1, 1 or 10 *μ*M oleuropein on the cell viability (O.D. 570 nm) of primary chondrocytes and hMSCs (Fig. 1b). We next examined hMSCs differentiation capacity in the presence of oleuropein (Fig. 1c, 1d and Supplementary Fig. 1). In accordance with previous results^52^, oleuropein-treated hMSCs showed significantly less in adipogenic differentiation detected by oil red O for lipid staining and by peroxisome proliferator-activated receptor (PPARγ) gene expression, whereas osteogenesis, examined by alizarin red staining of calcium deposits and by osteocalcin (OSTCN) gene expression, was significantly increased (Fig. 1c and Supplementary Fig. 1a and 1b). Furthermore, a 3D micromass culture system using chondrogenic medium supplemented with 10 *μ*M oleuropein or OE revealed an increase in ECM properties, reflecting a greater degree of chondrogenic differentiation within the 3D structure, with higher levels of Col2A1 deposited in the ECM and increased levels of aggrecan (ACAN) gene expression (Fig. 1d and Supplementary Fig. 1c). Remarkably decreased Cx43 gene expression was detected in hMSCs under the three differentiation conditions (adipogenesis, osteogenesis and chondrogenesis), this effect being more pronounced during chondrogenesis (Supplementary Fig 2a) and in concordance with a previous report^53^ (Fig. 1e and 1f). In this study, we have also observed changes in the Cx43 phosphorylation patterns at 7 and 14 days of chondrogenic differentiation by western blot (Fig. 1e), which can affect Cx43 stability and channel activity^54^. Different Cx43 phosphorylation patterns were also detected during adipogenesis and osteogenesis, suggesting differential regulation of Cx43 during hMSCs differentiation (Supplementary Fig. 2b). The treatment of hMSCs during chondrogenic differentiation with 10 *μ*M oleuropein for 14 days caused an additional 1.7 fold reduction in Cx43 gene expression (Fig. 1f). In concordance with these results, OACs’ differentiation (redifferentiation), for 7 days with chondrogenic medium supplemented with 10 *μ*M oleuropein, led to a significant decrease in the Cx43 protein levels without impact on Cx43 phosphorylation patterns (Fig. 1g).

**Figure 1.**
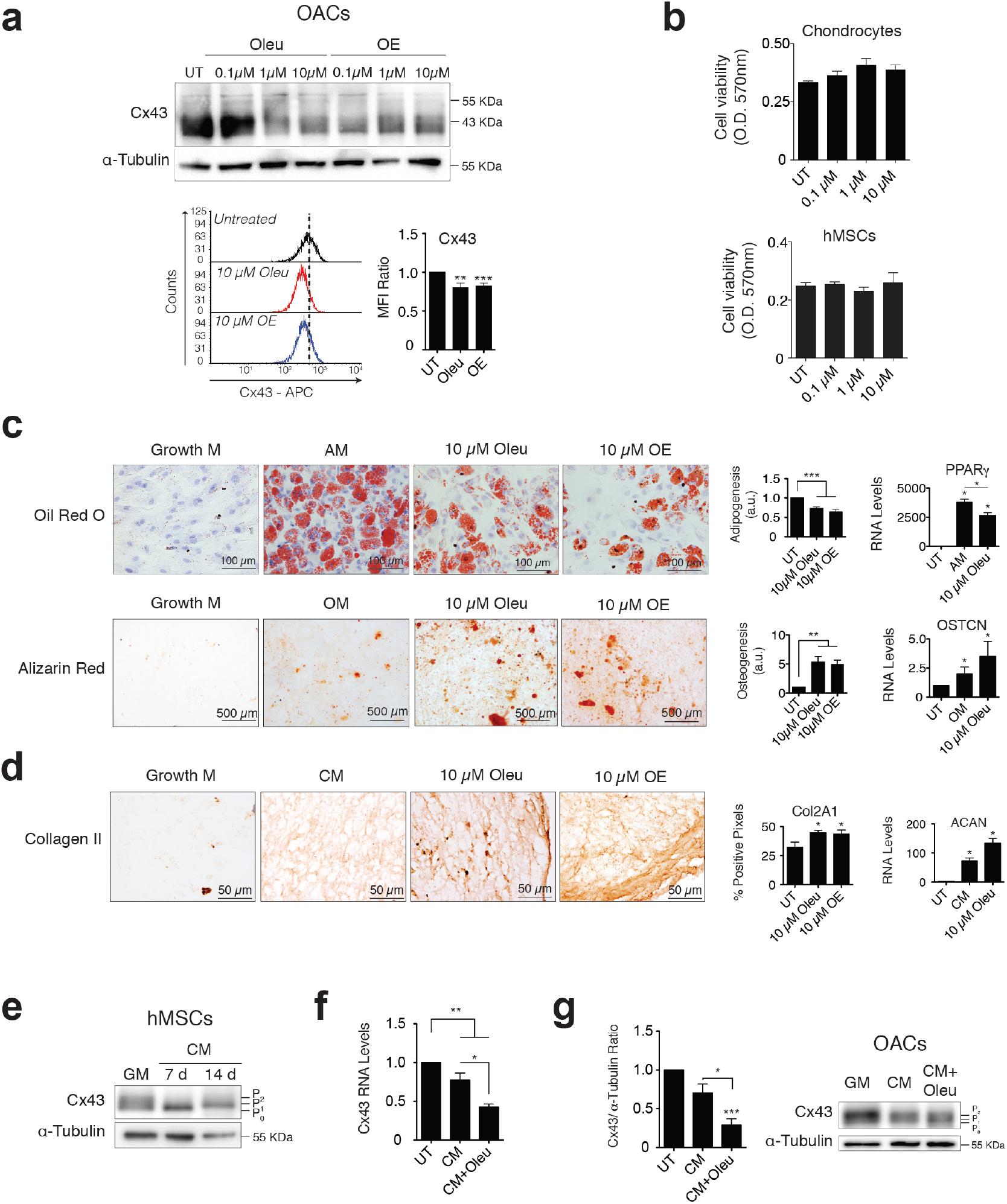
Downregulation of Cx43 during chondrogenesis improves differentiation towards chondrocytes. **(a)** Treatment of OACs with oleuropein (Oleu) or olive extract (OE) for 2 h significantly downregulates Cx43 protein (western blot and flow cytometry assays). The graph from the flow cytometry assays represents the median fluorescence intensity (n=9; mean±s.e.m.; \*\**P*<0.01, \*\*\**P*<0.0001; Mann–Whitney test). **(b)** Cell viability measured by MTT assay of hMSCs (n=5) and chondrocytes (n=7) exposed to different concentrations of oleuropein (Oleu) for 17 h (mean±s.e.m.). **(c)** Differentiation capacity of hMSCs grown in adipogenic (top, 21 days) or osteogenic (bottom, 21 days) medium supplemented with 10 *μ*M Oleu or 10 *μ*M OE. hMSCs cultured in growth medium were used as a control. Top, adipogenic evaluation by oil red O for lipid staining and by PPARγ gene expression (n=4-5; mean±s.e.m.; \**P*<0.05; Mann–Whitney test). Oil red O quantification represents the ratio of cells containing lipid deposits to the total number of cells (n=10-13 images from 2 independent experiments; mean±s.e.m.; \*\*\**P*<0.0001; Mann–Whitney test). Values were normalized to hMSCs differentiated in adipogenic medium without treatment (UT). Alizarin red staining was used to detect calcium deposits for osteogenic differentiation. Values were obtained by counting red pixels and normalized to those of hMSCs differentiated in osteogenic medium without treatment (UT) (n=4-6; mean±s.e.m.; \**P*<0.05, \*\**P*<0.01; Mann–Whitney test). OSTCN gene expression was measured to confirm osteogenic differentiation (n=4; mean±s.e.m.; \**P*<0.05; Mann–Whitney test). **(d)** Differentiation capacity of hMSCs grown in chondrogenic medium as micromasses for 30 days. Representative immunohistochemical images for Col2A1 indicate chondrogenic differentiation in micromasses. The quantification is shown on the right (n=4–6; mean±s.e.m.; \**P*<0.05; Mann–Whitney test). Chondrogenesis was also evaluated by ACAN gene expression quantification (n=3-4; mean±s.e.m.; \**P*<0.05; Mann–Whitney test). **(e)** Cx43 protein levels in hMSCs differentiated for 7 and 14 days in the presence of chondrogenic medium (CM) in comparison to untreated hMSCs cultured in normal growth medium (GM). **(f)** Cx43 RNA expression of hMSCs cultured for 14 days in the presence of chondrogenic medium (CM) alone or supplemented with 10 *μ*M oleuropein (Oleu). Data were normalized to HPRT-1 levels (n=5–6; mean±s.e.m. \**P*<0.05, \*\**P*<0.01; Mann–Whitney test). **(g)** Cx43 protein levels were analysed by western blot in OACs differentiated for 7 days in the presence of chondrogenic medium (CM), supplemented with 10 *μ*M oleuropein (Oleu). The graph represents the quantification from three independent experiments (mean±s.e.m.; \**P*<0.05, \*\*\**P*<0.0001; Student’s *t* test).

It is important to note that these results were obtained during differentiation of hMSCs under osteogenesis and chondrogenesis (Fig. 1c-f) and dedifferentiated chondrocytes from patients (OACs) in normal and in chondrogenic medium (Fig. 1a and Fig. 1g). However, the treatment of undifferentiated hMSCs cultured in basal growth medium with oleuropein or OE for 2 hours increased Cx43 levels (Supplementary Fig. 3a and 3b) and GJIC (Supplementary Fig. 3c), indicating that the effect of these olive derived polyphenols may be different depending on the cellular context.

### Downregulation of Cx43 activity by oleuropein downregulates the EMT transcription factor Twist-1 and enhances redifferentiation of OACs

Because oleuropein and the OE modulate Cx43 in OACs and the differentiation capacity of hMSCs, we expected that these compounds might modify de- and redifferentiation of OACs. Oleuropein treatment of primary chondrocytes isolated from patients with OA (OACs) decreased the Cx43 protein levels, as detected by western blot and flow cytometry (Fig. 1a and Fig 1g). We have previously found that intercellular coupling via GJ channels participates in the effects of Cx43 on the OAC phenotype^4^. Interestingly, oleuropein modulation of Cx43 significantly reduced GJIC in OACs, as detected by scrape loading/dye uptake assay and flow cytometry analysis (Fig. 2a), but not in healthy chondrocytes, with dye transfer through GJ channels being more reduced in OACs than in healthy chondrocytes (Fig. 2b). The decrease in Cx43 GJIC and protein levels were correlated with a significant reduction in the levels of the stemness markers CD105 and CD166 after 7 days in culture in presence of 10 *μ*M oleuropein as detected by western-blot and flow cytometry (Fig. 2c). This result was consistent with our previous observations, where CD105 and CD166 stemness markers were reduced when Cx43 was downregulated or when OACs were grown in chondrogenic medium to enhance redifferentiation for 14 days^4^. In fact, oleuropein effectively improved the phenotype of chondrocytes isolated from human OA cartilage grown in culture as a monolayer detected by Col2A1 levels and synthesis of proinflammatory mediators and MMP-3 (Fig. 2d and 2e). Treatment with 10 *μ*M oleuropein for 2 h was enough to reduce overall Cx43 positivity and to increase Col2A1 protein levels (Fig. 2d) and decrease interleukin 6 (IL-6), COX-2, IL-1β and MMP-3 gene expression (Fig. 2e).

**Figure 2.**
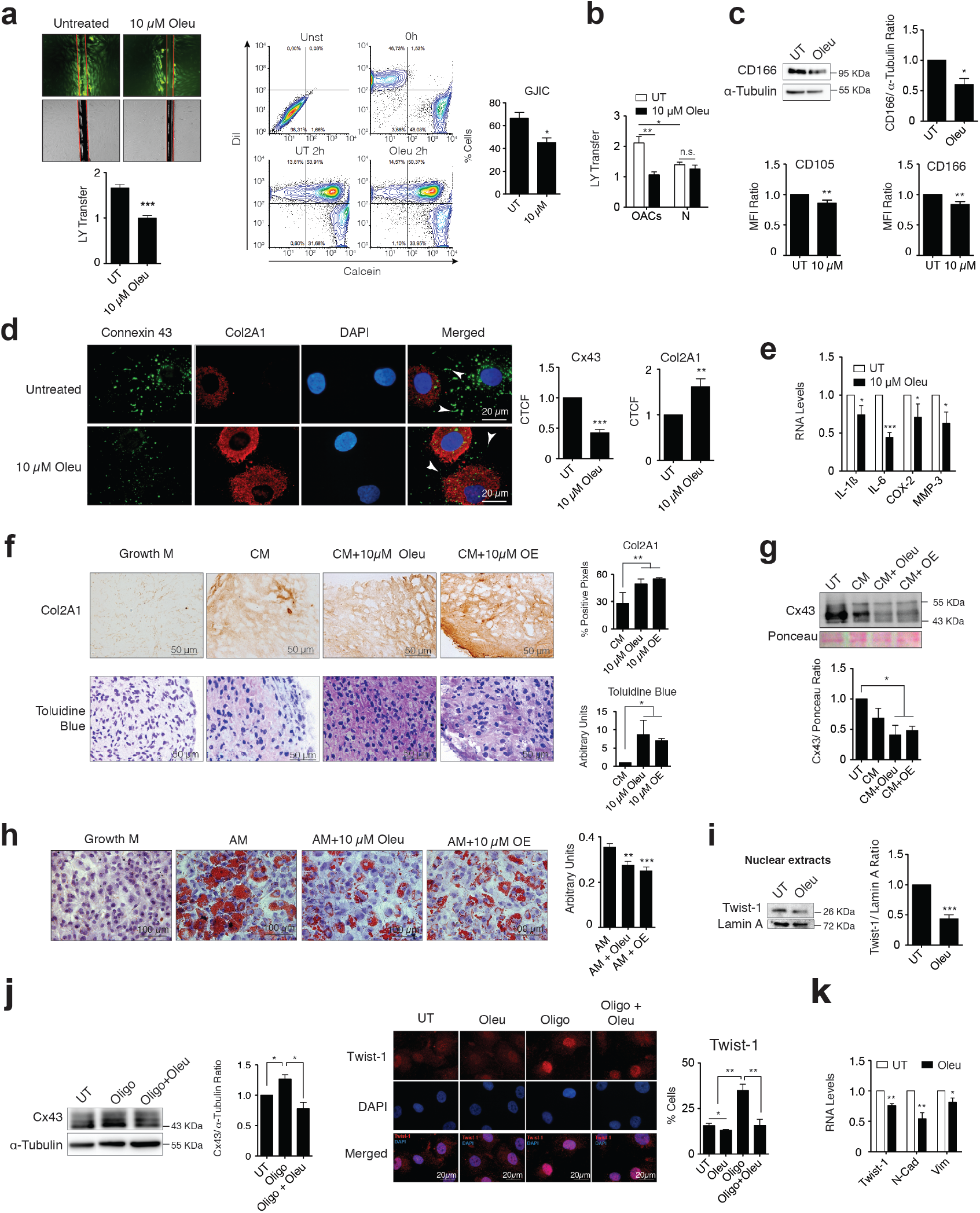
Downregulation of Cx43 by oleuropein restores chondrocyte redifferentiation in chondrocytes from patients with OA. **(a)** Oleuropein treatment significantly decreases GJIC evaluated by an SL/DT assay when OACs were exposed with this molecule for 2 h (top, n=6; mean±s.e.m.; \*\*\**P*<0.0001; Mann–Whitney test). The results were confirmed by calcein transfer and flow cytometry (n=4, mean±s.e.m.; \**P*<0.05; Mann–Whitney test). **(b)** Graph showing the effect of Oleu on GJIC when healthy chondrocytes (N) are exposed to 10 *μ*M Oleu compared with OACs (n=3; mean±s.e.m.; \**P*<0.05, \*\**P*<0.01; Mann–Whitney test). **(c)** OACs cultured for 7 days in DMEM 10% (UT) show reduced expression of the mesenchymal markers CD105 and CD166 when supplemented with 10 *μ*M Oleu. Protein levels were analysed by flow cytometry (n=7; mean±s.e.m.; \**P*<0.05, \*\**P*<0.01; Mann–Whitney test. In addition, CD166 levels were analysed by western blot (n=3; mean±s.e.m.; \**P*<0.05; Student’s *t* test). **(d)** Downregulation of Cx43 (green) by Oleu upregulates the main articular chondrocyte marker Col2A1 (red), as detected by immunofluorescence in OACs treated with Oleu or OE. Nuclei were stained with DAPI. Graphs represent the corrected total cell fluorescence (CTCF) measured with ImageJ with > 8 images per condition. Data represent mean±s.e.m (\*\**P*<0.01, \*\*\**P*<0.0001; Mann–Whitney test). **(e)** mRNA levels of IL-1β, IL-6, COX-2 and MMP-3 of OACs cultured in normal medium (UT, DMEM 10% FBS) exposed to 10 μM Oleu for 2 h (n=3–7; mean±s.e.m.; \**P*<0.05, \*\*\**P*<0.0001; Mann–Whitney test). **(f)** Immunohistochemistry (Col2A1) and toluidine blue staining (proteoglycan subunits) indicate significant enrichment in cartilage matrix components in OACs micromasses grown in 3D culture for 30 days in chondrogenic medium (CM) when supplemented with 10 *μ*M Oleu or OE (n=4–6 (Col2A1); n=6-7 (toluidine blue); mean±s.e.m.; \**P*<0.05; \*\**P*<0,01; Mann–Whitney test). **(g)** Cx43 levels detected by western blot (normalized to Ponceau staining) are reduced when OACs micromasses are exposed to chondrogenic medium (CM) supplemented with 10 *μ*M Oleu or OE for 21 days. (n=3 −4; mean±s.e.m.; \**P*<0.05; Mann–Whitney test). **(h)** Oil red staining showing reduced OACs dedifferentiation upon exposure to Oleu or OE in adipogenic medium. The graph represents the ratio of cells containing lipid deposits to the total number of cells (n=20 images; mean±s.e.m. \*\**P*<0.01, \*\*\**P*<0.0001; Mann–Whitney test). **(i)** Nuclear levels of Twist-1 were decreased in OACs cultured with 10 *μ*M Oleu for 2 hours. Lamin A was used as a loading control (n=3; mean±s.e.m, \*\*\**P*<0.0001, Student’s *t* test). **(j)** Twist-1 (red) activation measured by immunofluorescence to detect nuclear translocation when primary OACs were treated with 5 *μ*g/ml oligomycin for 1 h is partially abrogated by Cx43 downregulation and 10 *μ*M Oleu. Nuclei were stained with DAPI. The graph represents the percentage of cells with Twist-1 nuclear localization (n=4–5; mean±s.e.m, \**P*<0.05, \*\**P*<0.01; Mann–Whitney test). Scale bar, 20 μm. On the right, western blot (n=4) to show the effects of 1-h treatment with Oleu or oligomycin on Cx43 protein levels in primary OACs. Quantification is shown on the right (mean±s.e.m.; \**P*<0.05; Mann–Whitney test). **(k)** The RNA expression of the EMT markers Twist-1, N-Cadherin and Vimentin was upregulated in OACs treated with 10 *μ*M Oleu for 2 hours. Data were normalized to HPRT-1 levels (n= 5–6, mean±s.e.m, \**P*<0.05, \*\**P*<0.01; Mann–Whitney test).

Next, we sought to confirm whether oleuropein would target chondrocyte plasticity in 3D cultures. OACs were grown as a micromass pellet culture in chondrogenic medium supplemented with different concentrations of oleuropein (Fig. 2f). The enhancement of fully differentiated chondrocytes in chondrogenic medium improved the micromass matrix structure of OACs in 3D culture (Fig. 2f). Oleuropein and OE further decreased Cx43 levels (Fig. 2g) and further increased the deposition of proteoglycans detected by toluidine blue staining and Col2A1 detected by immunohistochemistry (Fig. 2f).

To further explore the effect of oleuropein on cell plasticity, OACs were grown in adipogenic medium in presence of 10 *μ*M oleuropein. Notably, oleuropein significantly decreased the adipogenic differentiation of OACs (Fig. 2h). However, oleuropein treatment increased the osteogenic differentiation of OACs when these cells were grown in osteogenic medium with 10 *μ*M oleuropein for 14 days (Supplementary Fig. 4). Consistent with this finding, previous studies have suggested a direct transformation of mature chondrocytes into osteoblasts and osteocytes^55–57^, reemphasizing the effect of oleuropein as a modulator of cell plasticity in chondrocytes, enhancing OACs redifferentiation to mature chondrocytes (and transdifferentiation into bone cells).

Upregulation of Cx43 in OA involves dedifferentiation via chondrocyte-to-mesenchymal transition (CMT) by Twist-1 activation, which was also reported in OA cartilage^4,58,59^. After oleuropein treatment, OACs showed a significant reduction in the nuclear localization of Twist-1 detected by western-blot of nuclear extracts (Fig. 2i). Furthermore, immunofluorescence assays showed that the nuclear localization of Twist-1 transcription factor in the presence of an arthritic insult (oligomycin), was attenuated when Cx43 protein levels were reduced by oleuropein treatment of OACs in monolayer culture (Fig. 2j). In addition to Twist-1, oleuropein treatment in OACs led to a significant reduction of the expression of mesenchymal and EMT markers N-cadherin and vimentin (Fig. 2k).

### Oleuropein modulates Cx43 gene promoter activity

Mechanisms that underlie the effects of oleuropein or the OE containing oleuropein and other small molecular polyphenols may be multifactorial. Yet, in chondrocytes decreased Cx43 protein levels were detected when OACs and the human chondrocyte cell line T/C-28a2 were treated with 10 *μ*M oleuropein (Fig. 1a and 3a). However, changes in Cx43 protein levels were not evident when T/C-28a2 transfected with a vector to overexpress Cx43 was treated with 10 *μ*M oleuropein (Fig. 3a, bottom), suggesting that oleuropein in chondrocytes may affect Cx43 gene promoter activity rather than protein stability. We thus measured whether oleuropein affected the activity of the Cx43 gene promoter using a real-time reporter system. Human T/C-28a2 chondrocytes were transfected with a firefly luciferase reporter vector containing the regulatory regions of the Cx43 promoter and incubated for 1 h with 10 *μ*M oleuropein and 5 *μ*g/ml of the mitochondrial inhibitor oligomycin, which inhibits ATP synthase by blocking its proton channel^60,61^ and enhances Cx43 gene expression (Fig. 2j, Fig 3b). In fact, this small molecule was previously used for *in vitro* and *in vivo* models of arthritis to induce inflammation and cartilage degradation^62^. Cx43 promoter activity (detected by luminescence) decreased after oleuropein treatment (Fig. 3b). Oligomycin increased the luminescence signals, and the effect of oligomycin was significantly attenuated in the presence of oleuropein (Fig. 3b), which was correlated with the Cx43 mRNA levels detected by RT-qPCR (Fig. 3b). The luminescence signals strongly correlated with the effects of oleuropein and oligomycin on protein levels detected by immunofluorescence (Fig. 3c) and western blot (Fig. 2j) when OACs were grown in monolayer culture. These results indicate that oleuropein affects Cx43 (gene, GJA1) promoter activity, thus downregulating Cx43 protein levels (Fig. 1a and Fig. 3a-c) and GJIC in OACs (Fig. 2b).

**Figure 3.**
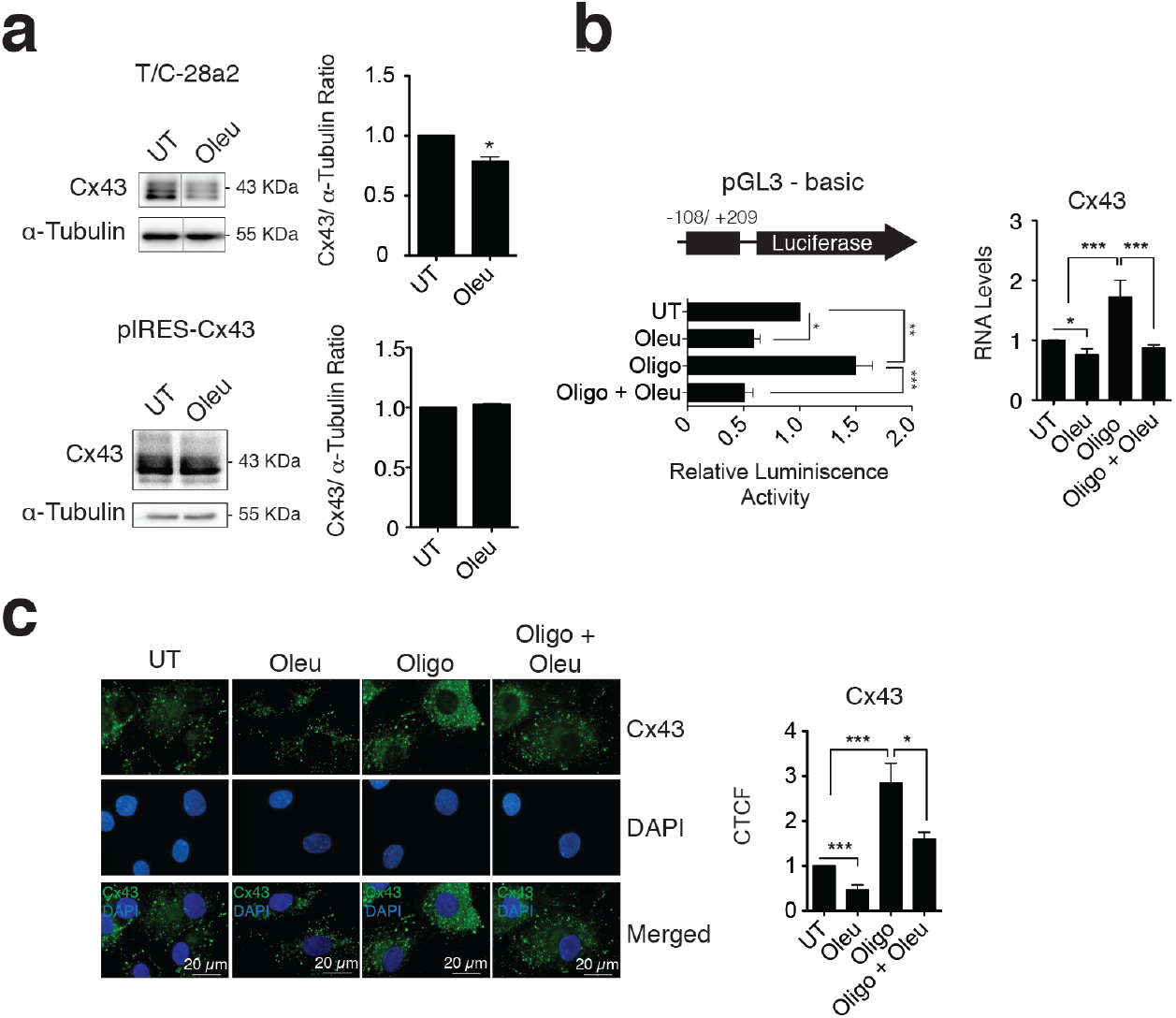
Oleuropein modulates the Cx43 promoter activity in chondrocytes. **(a)** Treatment with 10 *μ*M oleuropein (Oleu) for 2 h decreases Cx43 protein levels in T/C-28a2 cells (n=4), but this effect is not observed in the same cell line overexpressing Cx43 (pIRES-Cx43) (n=2, mean±s.e.m. \**P*<0.05; Mann–Whitney test). **(b)** Luciferase reporter assay indicating that Oleu inhibits Cx43 promoter activity. The graphs indicate the normalized luminescence activity in the T/C-28a2 chondrocyte cell line transfected with a pGL3-basic plasmid containing 300 base pairs of Cx43 promoter ligated to the luciferase gene. Cells were cultured in DMEM with 10% FBS (UT) and with 5 *μ*g/ml oligomycin or 10 *μ*M Oleu for 1 h as indicated (n=6; mean±s.e.m.; \*\**P*<0.01; Mann–Whitney test). On the right, Oleu reduces Cx43 gene expression and abolishes oligomycin-induced upregulation of Cx43 mRNA in OACs treated for 1 h following cell seeding (n=2-4; mean±s.e.m.; \**P*<0.05; \*\*\**p*< 0.0001; Student’s *t* test). **(c)** The effects of 1-h treatment with Oleu or oligomycin on Cx43 protein levels in primary OACs were confirmed by immunofluorescence (n=9, mean±s.e.m.; \**P*<0.05; \*\*\**p*< 0.0001; Mann–Whitney test).

### Oleuropein enhances elimination of senescent cells

Senescent cells through secretion of SASP promote dedifferentiation and reprogramming in neighbouring cells in the context of tissue injury^63,64^. OACs treated with 10 *μ*M oleuropein for 7 or 14 days in growth medium showed a significant reduction of senescent cells accumulated after 5 days of primary culture (Fig. 4a) and detected by beta-galactosidase activity measured by flow cytometry (Fig. 4a). Consistent with these models, Cx43 upregulation due to oligomycin insult (for 24 hours) significantly contributed to increase cellular senescence in OACs accumulated after 5 days in monolayer (Fig. 4b, left), whereas co-treatment with oleuropein significantly halted the accumulation of senescent cells, as measured by the quantitative flow cytometry analysis of beta-galactosidase activity (24 hours under treatments) (Fig. 4b, left) and beta-galactosidase staining (7 days under treatments) (Fig. 4b, right). Interestingly, increased levels of Cx43 were detected when the chondrocyte cell line T/C-28a2 was treated with 50 *μ*g/mL of bleomycin for 24 h hours and the exposure to oleuropein for 24 hours, after bleomycin treatment, reduced both the number of senescent cells detected by β-galactosidase activity and Cx43 protein levels detected by western-blot (Supplementary Fig. 5). In concordance with these results, the treatment of OACs with 10 *μ*M oleuropein for 2 hours led to a decrease in the levels of the senescence biomarker p16^INK4a^ (Fig. 4c) and p53 and p21 protein levels (Fig. 4d). Cellular senescence and the senescence-associated secretory phenotype (SASP) convert senescent cells into proinflammatory cells by secreting factors such as IL-6^65,66^. Oleuropein reduced the accumulation of senescent cells and attenuated oligomycin-induced SASP secretion detected by IL-6 and COX-2 gene expression in chondrocytes (Fig. 4e). This effect was also detected to a lesser extent for IL-1β. (Fig. 4e). The secretory activity of senescent cells (SASP), including IL-6 gene expression can be activated by the master regulator of SASP, NF-ĸB^67,68^. Immunofluorescence assays showed that NF-ĸB (p65) activation detected by nuclear translocation in OACs by TNFα was diminished when cells were exposed to 10 *μ*M oleuropein for 1 h (Fig. 4f). Oleuropein also protected from the increase of Cx43 under TNFα treatment (Fig. 4f, left). Further, NF-ĸB nuclear translocation was partially abolished in OACs after 5 days in primary culture and treated with 10 *μ*M oleuropein for only 2 hours and detected by western-blot (nuclear extracts) (Fig. 4g).

**Figure 4.**
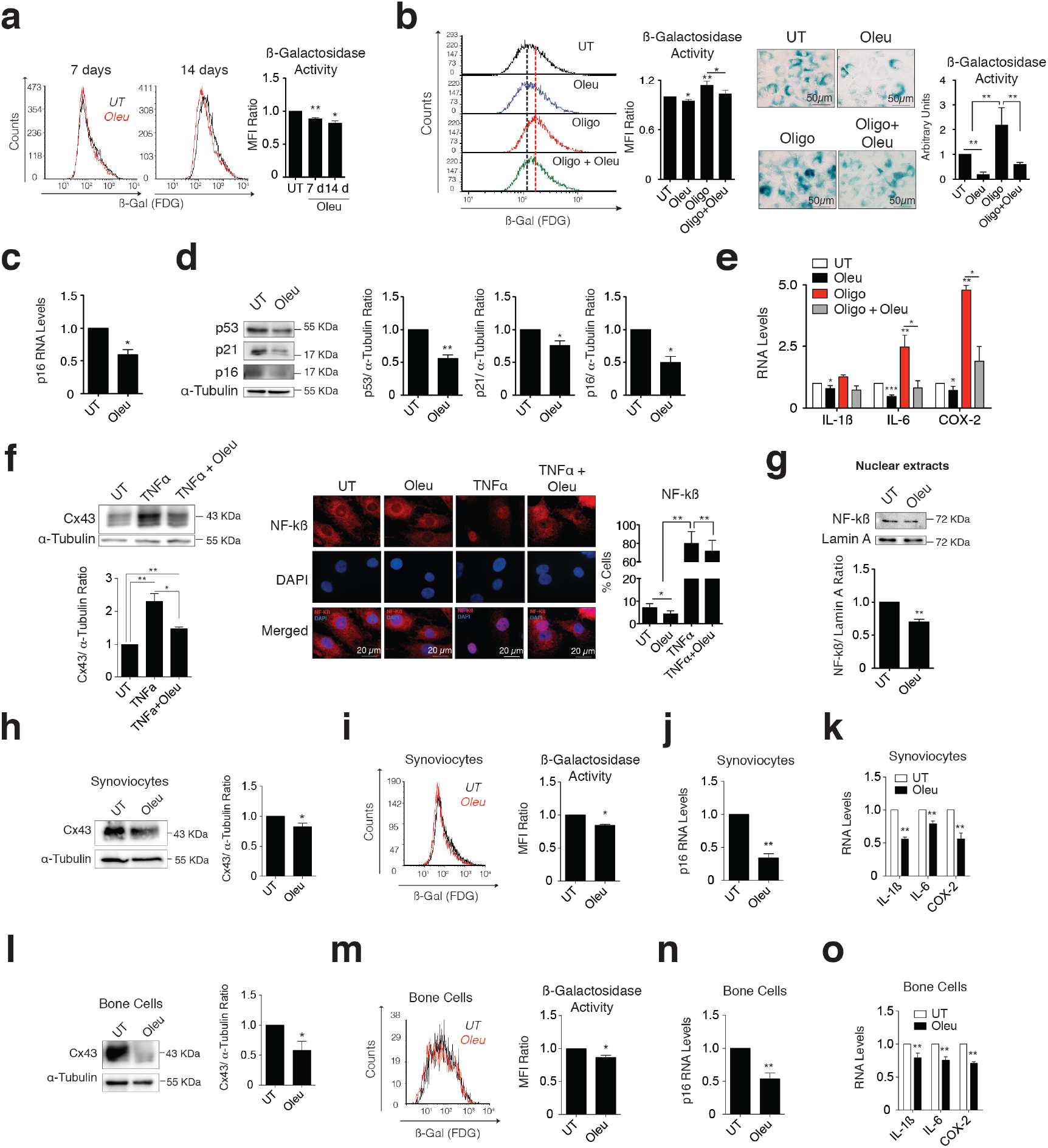
Cx43 downregulation by oleuropein decreased chondrocyte senescence. **(a)** Oleuropein (Oleu) treatment, which reduces Cx43 protein in OACs (See Fig. 1a), also reduces senescence as detected by SAβG and flow cytometry. Cells were treated with 10 *μ*M Oleu for 7 and 14 days (n=3–7; mean±s.e.m.; \**P*<0.05, \*\**P*<0.01; Mann–Whitney test). **(b)** Downregulation of Cx43 by Oleu treatment reduces senescence and abolishes oligomycin-induced senescence. The graphs show the comparative analysis of SAβG activity measured by flow cytometry of OACs exposed for 24 h to Oleu or oligomycin as indicated (n=5–6; mean±s.e.m.; \**P*<0.05, \*\**P*<0.01; Mann–Whitney test). Right, to further confirm these results, SAβG activity, determined by X-Gal cleavage and cell staining (blue), was evaluated by microscopy in OACs treated for 7 days with these drugs. The images are representative from three independent experiments and quantification is shown on the right (mean±s.e.m.; \**P*<0.05, \*\**P*<0.01, \*\*\**P*<0.0001; Mann–Whitney test). **(c)** p16 mRNA expression of OACs treated with 10 *μ*M Oleu for 2 h. Data were normalized to HPRT-1 levels (n= 5, mean±s.e.m, \**P*<0.05; Mann–Whitney test). **(d)** Western blot of p53, p21 and p16 levels in OACs treated with 10 *μ*M oleuropein (Oleu) for 2 h. α-tubulin was used as a loading control (n=2-3; mean±s.e.m.; \**P*<0.05, \*\**P*<0.01; Student’s *t* test). **(e)** Downregulation of Cx43 by Oleu attenuates the IL-6 and COX-2 upregulation when OACs are exposed to oligomycin for 1 h (n=3-9; mean±s.e.m.; \**P*<0.05, \*\**P*<0.01, \*\*\**P*<0.0001; Mann–Whitney test). **(f)** Western blot (n=3) shows the effect of Oleu and TNFa treatments on Cx43 protein levels in primary OACs (mean±s.e.m.; \**P*<0.05, \*\**P*<0.01; Student’s *t* test). On the right, NF-ĸB (red) detected by immunofluorescence in OACs treated with TNFa to induce NF-ĸB nuclear translocation and activation. This effect is partially abolished by 1-h Oleu treatment. Nuclei were stained with DAPI. The graph represents the cell percentage with nuclear NF-ĸB staining (n=5; mean±s.e.m.; \**P*<0.05; Mann–Whitney test). (Pending WB) **(g)** Nuclear levels of NF-kβ were decreased in OACs cultured with 10 *μ*M Oleu for 2 hours. Lamin A was used as a loading control (n=3; mean±s.e.m, \*\**P*<0.01, Student’s *t* test). **(h)** Cx43 protein levels analysed by western blot in synoviocytes treated with oleuropein (Oleu) for 2 hours. a-tubulin was used as a loading control (n=7; mean±s.e.m.; \**P*<0.05; Mann–Whitney test). **(i)** Treatment of synoviocytes with 10 *μ*M of oleuropein (Oleu) for 7 days reduces cell senescence, as detected by SAβG and flow cytometry (n=4; mean±s.e.m.; \**P*<0.05; Mann–Whitney test). **(j)** p16 mRNA levels of synoviocytes treated with 10 *μ*M Oleu for 2 h. Data were normalized to HPRT-1 levels (n= 4, mean±s.e.m, \*\**P*<0.01; Student’s *t* test). **(k)** mRNA levels of IL-1β, IL-6 and COX-2 of synoviocytes cultured in normal medium (UT, DMEM 10% FBS) exposed to 10 *μ*M Oleu for 2 h. Data were normalized to HPRT-1 levels (n=4; mean±s.e.m.; \*\**P*<0.01; Student’s *t* test). **(l)** Cx43 protein levels analysed by western blot in bone cells treated with oleuropein (Oleu) for 2 hours. α-tubulin was used as a loading control (n=4; mean±s.e.m.; \**P*<0.05; Mann–Whitney test). **(m)** 10 *μ*M of oleuropein (Oleu) treatment for 7 days reduces senescence levels in bone cells as detected by SAβG and flow cytometry (n=3; mean±s.e.m.; \**P*<0.05; Student’s *t* test). **(n)** p16 mRNA expression of bone cells treated with 10 *μ*M Oleu for 2 h. Data were normalized to HPRT-1 levels (n= 4, mean±s.e.m, \*\**P*<0.01; Student’s *t* test). **(o)** mRNA levels of IL-1β, IL-6 and COX-2 of bone cells cultured in normal medium (UT, DMEM 10% FBS) exposed to 10 *μ*M Oleu for 2 h. Data were normalized to HPRT-1 levels (n=3-4; mean±s.e.m.; \*\**P*<0.01; Student’s *t* test).

To further test the senolytic activity of oleuropein, synovial and bone cells isolated from synovium and spongy bone from joints from OA patients were treated with 10 *μ*M oleuropein (Fig. 4h-o). We observed decreased Cx43 protein levels after only 2 hours under oleuropein exposure (Fig. 4h and 4l) together with a significant reduction in senescent cells accumulated after 5 days in primary culture and detected by β-galactosidase activity (Fig. 4i and 4m) and confirmed by RT-qPCR to test the expression of the senescence biomarker p16^INK4a^ (Fig. 4j and 4n) and the synthesis of the SASP factors IL1-β, COX-2 and IL-6 (Fig. 4k and 4o).

## Discussion

Chondrocyte phenotypic changes are common response to insults and during OA progression^4^. Previous data reported by our group demonstrate that downregulation of Cx43 improves cell phenotype protecting chondrocytes from dedifferentiation (via EMT transcription factors such as Twist-1) and senescence (via p53 activation)^4^. Although oleuropein was reported to protect from OA progression^69–73^, there was no solid evidence about the underlying molecular mechanisms involved in this effect. Our results are consistent with previous studies showing the protective effect of oleuropein and olive derived polyphenols in OA^69,71^ and that Cx43 downregulation have significant effects on chondrocyte phenotype^4,16,30^. In our study, we show that oleuropein modulates the activity of the Cx43 gene promoter, reducing Cx43 and GJIC in OACs and hMSCs. Indeed, our data indicate that downregulation of Cx43 by oleuropein in OACs in 3D and monolayer culture improves cell phenotype by protecting chondrocytes from the EMT transcription factor Twist-1 activation under experimental insults and from accumulation of senescent cells (Fig. 2 and Fig. 4). This is the first study that demonstrated one of the potential mechanisms of oleuropein in OACs, synovial and bone cells (Fig 4) from patients and in hMSCs (Fig. 1), with important applications in regenerative medicine.

Experimental, clinical and epidemiological data have pointed out the beneficial effects of polyphenols in wound healing in multiple ageing-related disorders^74–77^, making these compounds effective candidates in the context of preventive nutrition and for the development of new therapeutic strategies^77–81^. Oleuropein is the most abundant polyphenol in the leaves and fruit of the olive plant (e.g. *Olea europaea* L.) and is a potent antioxidant agent with anti-tumour and anti-inflammatory properties^82,83^. The mechanism of action this polyphenol is under study^82^, and show that oleuropein and its major metabolite hydroxytyrosol have antioxidant activity by inhibition and/or scavenging of ROS^84,85^, which has been suggested to reduce NF-kB activation^86–89^. Other mechanistic studies implicate nitric oxide (NO) production^89–92^ or autophagy and inhibition of the mammalian target of rapamycin (mTOR)^93–95^. Here, we show that oleuropein directly regulates Cx43 gene promoter activity leading to downregulation of GJIC (Fig. 2a and Fig. 3b). Interestingly, in hMSCs oleuropein reduces Cx43 and GJIC only under differentiation processes and increases Cx43 in undifferentiated hMSCs (Fig 1e, 1f, Supplementary Fig. 2 and Supplementary Fig. 3). These results indicate that the activity of this molecule in the Cx43 gene promoter depends on the context. In fact, a gene expression profiling study has suggested that oleuropein affects the expression of multiple genes involved in oxidative stress, inflammation, fibrosis, cell proliferation and differentiation among others^96^, which implies that the mechanisms that underlie the beneficial effects of this molecule may be multifactorial and depend on the conditions^96^. These observations emphasize the importance of Cx43-mediated effects of oleuropein specifically during the differentiation process. Also, our results show that the effects of oleuropein are often equal or even smaller than those of olive extract that contains 41.5% of oleuropein at equivalent doses. This observation suggests that other polyphenols or compounds, such as the hydroxytyrosol, may synergize with oleuropein activity in chondrocytes^97,98^. On the other hand, the treatment of hMSCs with oleuropein or OE, in chondrogenic, osteogenic or adipogenic differentiation medium leads to downregulation of Cx43 and GJIC, enhancing osteogenesis and chondrogenesis, but reducing adipogenesis. This differential sensitivity of hMSCs to oleuropein and OE may have potential applications in preventive and regenerative medicine in other bone and cartilage disorders e.g. by reducing ectopic adipocyte accumulation in bone marrow cavities and improving osteogenesis during bone repair^99–101^.

Our data show decreased Cx43 levels with decreased GJ activity in the presence of oleuropein in OACs and in differentiating hMSCs, but increased Cx43 levels with increased GJIC in undifferentiated hMSCs, suggesting that the effect of this molecule on GJIC depends on its effect on Cx43 levels (or subcellular localization). In fact, phosphorylation of Cx43 affects protein stability and GJIC activity^54,102,103^ and we have detected changes in Cx43 phosphorylation pattern during hMSCs differentiation (chondrogenesis, osteogenesis and adipogenesis) but not under oleuropein treatment (in hMSCs and OACs) indicating that oleuropein affects Cx43 levels more than protein or gap junction plaque stability or regulation of the activity of these intercellular channels. Further, Cx43 channel-independent activities involve its signalling hub’s ability to recruit proteins to the membrane^24,104–106^ or its ability to control gene transcription by nuclear translocation and binding to RNA polymerase II and gene promoters^107^. We have recently reported that overactivity of Cx43 in OACs compromise the ability of these cells to redifferentiate by maintaining the stem-like state enhanced by Twist-1 activity and tissue remodelling and proinflammatory agents such as IL-1β^4^. Here, we show that oleuropein restores chondrocyte phenotype detected by reduced levels of the stem-markers CD105, CD166, N-cad and vimentin (Fig. 2c, 2k). The effect of oleuropein in chondrocyte plasticity correlated with activation of redifferentiation via downregulation of Cx43 and Twist-1 (Fig.2i, 2j, 2k), leading to increased levels of proteoglycans and Col2A1 together with a decrease in the levels of inflammatory cytokines and metalloproteinases such as IL-1β and MMP-3 (Fig. 2d, 2e, 2f).

In cell culture and in osteoarthritic cartilage, OACs undergo dedifferentiation and senescence^4,108,109^. Elimination of senescent cells *in vivo* using the senolytic drug UBX0101 has been demonstrated to improve cartilage regeneration after articular joint injury in mice^3^. Here we show that oleuropein reduces cellular senescence in OACs, synovial and bone cells in monolayer and protects from accumulation of senescent cells under an arthritic insult (Fig. 4a, 4b, 4i and 4m). NF-kB has been shown to cooperatively regulate the inflammatory components of the SASP, together with other factors^68,110–112^. In this study the reduction of senescence is accompanied by reduced activity of the NF-kB transcription factor and therefore reduced synthesis of SASP including those encoding IL-6, IL-1β and COX-2 (Fig. 4e). Notably, these components enhance inflammation, senescence and activate dedifferentiation and cellular reprogramming of nearby non-senescent cells (e.g. via IL6)^63,64^, contributing to the stem-like state of chondrocytes in OA and to the accumulation of senescent cells. Accordingly, we have previously reported that upregulation of Cx43 leads to p53/p16 upregulation and senescence^4^. Using the T/C-28a2 human chondrocyte cell line with a Cx43 overexpression vector and a CRISPR/Cas9-mediated heterozygous Cx43 gene knockout cell line we have demonstrated that Cx43 is an upstream effector of both senescence (involving p53 and p16 pathways) and NF-kB activation (SASP synthesis)^4^. However IL-1 signalling was also shown to be an upstream effector of NF-kB^113–115^. IL-1 gene expression is activated by Cx43^4,29^ and vice versa, providing another level of positive feedback loop involved in cartilage degradation in OA.

Cellular reprogramming, dedifferentiation via EMT and senescence play active roles during tissue regeneration^116^. These processes are activated in response to damage and involve dedifferentiation, reprogramming, inflammation and ECM remodelling and resolve with redifferentiation and elimination of senescent cells^116^. Accumulation of dedifferentiated (stem-like cells) and senescent cells leads to impaired tissue regeneration and fibrosis with loss of tissue function^117,118^ (Fig. 5). Understanding and manipulating the complex Cx43 signalling appears to be very challenging but would expand our opportunities for modulating wound-healing related disorders such as OA. So far, our results indicate that molecules that reduce Cx43 levels in OA such as oleuropein will potentially restore cartilage repair in OA by activating chondrocyte redifferentiation and by eliminating senescent cells, contributing to creating a regeneration-permissive environment in OA patients.

**Figure 5.**
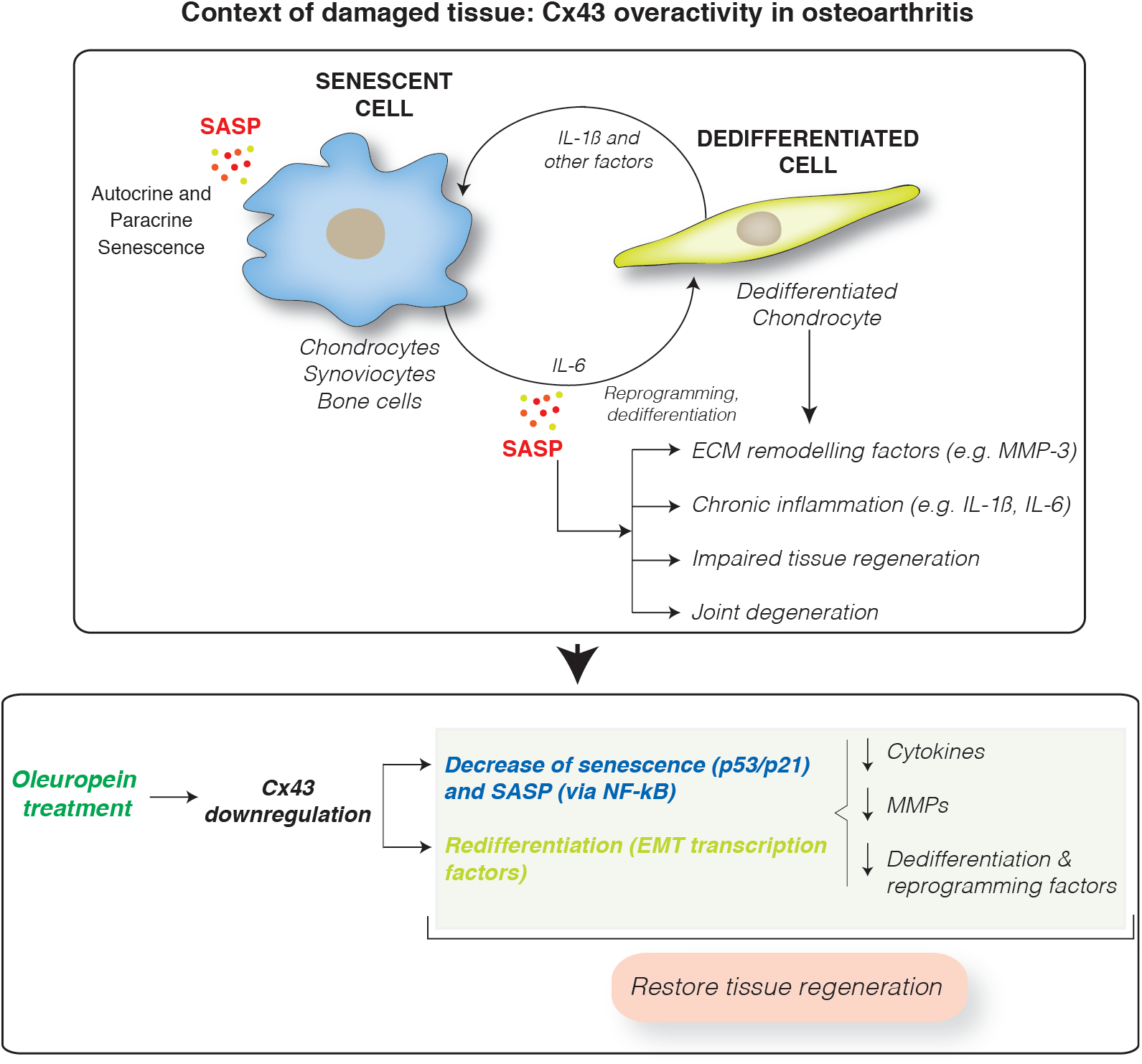
Cx43 overexpression leads to accumulation of dedifferentiated and senescent cells involved in disease progression in OA patients. These phenotypic changes results in the synthesis of ECM remodelling factors involved in tissue degradation (MMPs) and proinflammatory factors, such as IL-1β and IL-6, which facilitate the dedifferentiation and reprogramming of neighbouring cells. These factors may further spread senescence and dedifferentiation to surrounding tissues contributing to joint degeneration. Downregulation of Cx43 by oleuropein treatment contributes to the elimination of senescent cells and redifferentiation of osteoarthritic chondrocytes into fully differentiated cells, able to support the ECM composition and restoring the regenerative capacity of the tissue.

Consistent with previous studies, oleuropein shows a potent senolytic and anti-inflammatory effect by reducing IL-1β, IL-6 and COX-2 gene expression and NF-kB activation^119^. Our study is the first to demonstrate the effect of this polyphenol on Cx43 activity and senescence. Our results show that oleuropein downregulates Cx43 in OACs, enhancing chondrocyte redifferentiation, together with a decrease in cellular senescence and SASP activated by NF-kB (Fig. 5). Besides, modulation of Cx43 by oleuropein shifts the hMSCs differentiation capacity towards osteogenic and chondrogenic lineages, while decreasing adipogenic differentiation. These effects may be useful in cell therapy approaches that aim to promote cartilage and bone regeneration because the osteogenesis/adipogenesis switch has been associated with different bone disorders characterized by reduced bone formation and increased bone marrow fat accumulation^99,100^. Altogether, these findings indicate that oleuropein controls Cx43 gene expression and acts as a potent senolytic drug that may serve as a potential agent to improve both the effectiveness of stem cell therapy and cartilage and joint regeneration in patients to stop OA progression, by restoring tissue regeneration.

## Experimental procedures

### Tissue collection, processing and cell culture

Collection and processing of cartilage from human knees and femoral heads from adult donors undergoing joint surgery were performed as previously described^28^. The study was conducted with the approval of the institutional ethics committee (C.0003333, 2012/094 & 2015/029) after the acquisition of the signed informed consents. Samples from healthy, moderate OA (grades I–II) and high OA (grades III–IV) groups were selected for this study, which were classified according to their medical record data and histological analysis, as previously described^27^. 25 human samples were used for this study. Each experiment was conducted with at least three different patients/samples. Primary chondrocytes, isolated from fresh cartilage as previously described^28^, were cultured in Dulbecco’s Modified Eagle’s medium (DMEM) supplemented with 100 U/ml penicillin, 100 *μ*g/ml streptomycin and 10% foetal bovine serum (FBS), all from Gibco, Thermo Fisher Scientific. The chondrocyte cell line T/C-28a2, kindly donated by Dr. Goldring from the Hospital for Special Surgery (New York, USA), was cultured in DMEM supplemented with 10% FBS, 100 U/ml penicillin and 100 *μ*g/ml streptomycin. Human mesenchymal stem cells were obtained with signed informed consent from bone marrow donors *(Hospital Universitario Reina Sofía*, Córdoba, Spain) and from subcutaneous inguinal fat from healthy individuals *(Hospital Universitario Ramón y Cajal*, Madrid, Spain). hMSCs were cultured in α-minimum essential medium (α-MEM; Lonza) supplemented with 10% FBS, 100 U/ml penicillin, 100 *μ*g/ml streptomycin, 2 mM GlutaMax (Gibco, Thermo Fisher Scientific) and 1 ng/ml recombinant human fibroblast growth factor-2 (rhFGF-2; Immunotools) or in MesenPRO RS™ Medium supplemented with 100 U/ml penicillin and 100 *μ*g/ml streptomycin.

For cell treatments, oleuropein with ≥ 90% purity was purchased from Extrasynthese (0204) (Lyon, France) and OE was donated by the Clinical Management Unit of Endocrinology and Nutrition (IMIBIC, Córdoba, Spain). This extract contains 41.5% oleuropein and was dissolved in culture medium to a stock concentration of 13 mg/ml (equivalent to 100 *μ*M oleuropein). Both compounds were dissolved in the cell culture medium and added to the cells for short-term (1–2 h) or long-term (7–21 days) treatments. Cell insults were performed with either 5 *μ*g/ml oligomycin (Sigma-Aldrich, O4876) or 10 ng/mL TNFα (Immunotools, 11343013), for 1 h.

### Cell viability assay

Cells in 96-well culture plates were treated with 0.1 μM, 1 μM, 10 *μ*M and 10 mM oleuropein for 17 h. Drug cytotoxicity was evaluated by the colorimetric MTT (3-(4,5-dimethylthiazol-2-yl)-2,5-diphenyltetrazolium bromide) assay (Cell Proliferation Kit I, Roche) with a NanoQuant microplate reader (Tecan Trading AG, Switzerland) at 570 nm.

### Adipogenic differentiation

Adipogenic differentiation was performed as previously described^4^. hMSCs or chondrocytes at a 80 −100% confluence were incubated with adipogenic medium (hMSC Adipogenic Differentiation Bullekit™, Lonza) for 21 days.

The medium was changed every 2 – 3 days, with the addition of 0.1, 1 or 10 *μ*M oleuropein or OE when necessary. Adipogenic differentiation was evaluated by the RNA expression of PPARγ (5’-GCGATTCCTTC ACTGATACACTG-3’, 5’-GAGTGGGAGTGGTCTTCCATTAC-3’) and by oil red O staining with a freshly prepared 60% (v/v) oil red O solution. Lipid droplet-containing cells were analysed with the ImageJ software (version 1.48) and adipogenic differentiation was evaluated as the ratio of positive droplet-containing cells to the total number of cells.

### Osteogenic differentiation

For osteogenic differentiation, cells were differentiated for 21 days with osteogenic medium (StemPro^®^ Osteogenesis Differentiation Kit, Gibco, Thermo Fisher Scientific), changing the medium every 2 – 3 days with the addition of 0.1, 1 or 10 *μ*M oleuropein or OE. Osteogenesis was evaluated by the RNA expression of OSTCN (5’-CCATGAGAGCCCTCACACTCC-3’, 5’-GGTCAGCCAACTCGTCACAGTC-3’) and by alizarin red S staining with a 2% (w/v) alizarin red S solution (Sigma-Aldrich). Slides were imaged in an Olympus BX61 microscope, and red positivity was analysed using ImageJ software (version 1.48).

### Chondrogenic differentiation

hMSCs or chondrocytes were differentiated with chondrogenic medium (StemPro^®^ Chondrogenesis Differentiation Kit, Gibco, Thermo Fisher Scientific) as micromasses for 30 days in the presence of chondrogenic medium, with or without oleuropein or OE addition. Micromasses were embedded in Tissue-Tek^®^ OCT™ (Sakura) and chondrogenesis was evaluated with ECM-specific stains (toluidine blue, safranin O-fast green) and by ACAN RNA expression (5’-CAGAACAACTCGGGGAACAT-3’, 5’-GCACAATTGGAACCCTGACT-3’).

### Scrape loading/dye transfer (SL/DT) assay

In order to evaluate GJIC, a SL/DT assay was performed as previously described^4,28^. Confluent cells were cultured for 16 h of in DMEM without FBS supplementation, and then oleuropein or olive leaf extract (OE) was added to the medium for 2 h. Next, a 0.4% (w/v) solution of lucifer yellow (LY) (Cell Projects Ltd©, Kent, UK) in PBS was loaded and two distant scrapes were made across the culture plate. The LY solution was absorbed by the damaged cells, which were allowed to transfer the dye for 3 min at 37°C. LY transfer was evaluated in a Nikon Eclipse Ti fluorescent microscope, and the number of dye-positive cells (LY transfer) from the cut site to the farthest detectable uptake of LY reflects the GJ connectivity between cells. The score was calculated as previously reported^120,121^.

### GJ connectivity by flow cytometry

Equal numbers of cells were incubated for 1 h at 37 °C with either 1 *μ*M of the membrane dye DiI (Invitrogen, Thermo Fisher Scientific) or 1 *μ*M calcein-AM (Invitrogen, Thermo Fisher Scientific), a cell-permeable dye that is retained in the cytosol and can be transferred by GJs.. Then, cells were washed 3 times with PBS and co-cultured for 2 h in a 1:2 ratio (calcein-donors:DiI-recipient cells). The percentage of Dil^+^Calc^+^ cells was analysed by flow cytometry and compared with that of unlabelled cells and to a 1:1 mixture of labelled cells mixed immediately before the analysis (Time = 0 h).

### Western blot

For protein analysis, cells were lysed in ice-cold lysis buffer (150 mM NaCl, 50 mM Tris-HCl pH 7.5, 5 mM EDTA pH 8, 0.5% v/v Nonidet P-40, 0.1% (w/v) SDS, 0.5% (v/v) sarkosyl) supplemented with 5 *μ*g/ml protease inhibitor cocktail and 1 mM phenylmethylsulfonyl fluoride (PMSF; Sigma-Aldrich). Nuclear protein isolation was performed with the NE-PER™ kit (Thermo Fisher Scientific), according to the manufacturer’s instructions. Proteins were separated on a 10% SDS-PAGE and transferred to a polyvinylidene fluoride (PVDF) membrane (Millipore Co., Bedford, MA). Primary antibody incubation was performed overnight at 4°C, and HRP-secondary antibody incubation was performed at RT for 1 h. Signal developing was performed with the Pierce™ ECL Western Blotting Substrate in either a LAS-300 Imager (Fujifilm) or an Amersham Imager 600 (GE Healthcare). The following primary antibodies were used: α-tubulin (Sigma-Aldrich, T9026), Cx43 (Sigma-Aldrich, C6219), Twist-1 (Santa Cruz Biotechnology, sc-81417), CD166 (Santa Cruz Biotechnology, sc-74558), p16^INK4a^ (Abcam, ab108349), p21 (Santa Cruz Biotechnology, sc-6246), p53 (Santa Cruz Biotechnology, sc-126), NF-ĸB (Santa Cruz Biotechnology, sc-8008) and lamin A (Santa Cruz Biotechnology, sc-20680).

### Antigen expression analysis by flow cytometry

Paraformaldehyde-fixed cells were incubated with phycoerythrin (PE)-conjugated anti-human CD105 (Immunostep, 105PE-100T) or allophycocyanin (APC)-conjugated anti-human CD166 (Immunostep, 1399990314), for 30 min at 4°C in the dark, as previously reported^4^ (Supplementary table 1). Intracellular Cx43 protein was detected in paraformaldehyde-fixed cells, which were permeabilized with methanol for 30 min at 4°C. Finally, cells were incubated with APC-conjugated anti-human Cx43 antibody (R&D Systems, FAB7737A; Supplementary table 1) for 30 min at 4°C in the dark.

### Flow cytometry analysis

Between 10.000 – 20.000 events were collected on a BD FACSCalibur™ (Becton Dickinson) flow cytometer with the CellQuest™ Pro software. Cell debris was discriminated by the forward scatter (FSC) and side scatter (SSC) properties of the cells. Data were analysed with FCS Express 6 Flow software (De Novo Software). The level of positive staining was expressed as the median fluorescence intensity (MFI), with unlabelled cells as negative controls. Gates were placed based on single-labelled controls and by establishing 0.1% as the cutoff point.

### Senescence-associated β-galactosidase activity

Flow cytometry analysis of SAβG activity with the fluorogenic β-galactosidase substrate fluorescein di-β-D-galactopyranoside (FDG; Invitrogen, Thermo Fisher Scientific) was performed as previously described^4^. Briefly, harvested cells were incubated with pre-warmed 2 mM FDG for 3 min at 37 °C, and fluorescein positivity was analysed on a BD FACSCalibur™ (Becton Dickinson) flow cytometer. SAβG activity was also measured with the Senescence Cells Histochemical Staining Kit (Sigma-Aldrich) according to the manufacturer’s protocol.

### Immunofluorescence

2% paraformaldehyde-fixed cells were incubated for 10 min with 0.1 M glycine (Sigma-Aldrich). Membrane permeabilization was performed with 0.2% Triton X-100 (Sigma-Aldrich) in PBS, 10 min at RT, followed by a 30-min incubation with 1% BSA and 0.1% Tween 20 in PBS (PBST). Primary and secondary antibodies were diluted in 1% BSA in PBST and incubated for 1 h at RT. Nuclei were stained with 1 *μ*g/ml 4’,6-diamidino-2-phenylindole dihydrochloride (DAPI; Sigma-Aldrich) in the dark for 4 min at RT. Slides were mounted with glycergel aqueous mounting medium (Dako). The following primary antibodies were used: anti-Cx43 (Sigma-Aldrich, C6129), anti-collagen II (Invitrogen, Thermo Fisher Scientific, MA5-12789), anti-Twist-1 (sc-81417) and anti-NF-ĸB (sc-8008) from Santa Cruz Biotechnology. Goat anti-rabbit FITC-conjugated (F-2765, Invitrogen, Thermo Fisher Scientific) and goat anti-mouse Alexa 594-conjugated (A-11032, Invitrogen, Thermo Fisher Scientific) secondary antibodies were used. Fluorescence was analysed by using ImageJ software version 1.48 and is shown as the corrected total cell fluorescence (CTCF).

### Immunohistochemistry

Micromass sections were fixed with acetone for 10 min, PBS-washed and nonspecific endogenous peroxidase activity was quenched with a 3% (v/v) hydrogen peroxidase solution (Roche) for 10 min. Primary antibody was incubated for 1 h at RT, and then slides were washed with PBS and incubated for 10 min with OptiView HQ Universal Linker (Roche) at RT. After three additional PBS-washes, slides were incubated with OptiView HRP Multimer (Roche) for 8 min at RT. Peroxidase activity was developed using a 0.02 % hydrogen peroxidase solution and 0.1 % DAB solution. Sections were then washed in distilled water and, in some cases, counterstained with Gill’s hematoxylin (Merck Millipore). Finally, slides were dehydrated with alcohol, cleared with xylene (PanReac AppliChem) and mounted with DePeX (BDH Gun^®^, VWR).

### Immunohistochemistry analysis

For immunohistochemistry analysis quantification, an in-house developed MATLAB program was used. This program first splits RGB images into single channels and applies a 3-level automatic thresholding to the green channel using Otsu’s method. Then, it segments the images into 4 discrete classes using the threshold levels. As a final step, the total pixels are obtained from the segmented images belonging to the 3 darker classes by discarding the lightest level in the image (background).

### Quantitative PCR

TRIzol™ reagent (Invitrogen, Thermo Fisher Scientific) was used to isolate total RNA, according to the manufacturer’s instructions. Degradation of DNA from samples was ensured by treatment with RNase-free DNase (Invitrogen, Thermo Fisher Scientific). 1 *μ*g of total RNA per reaction was used to synthesize cDNA with the SuperScript^®^ VILO™ cDNA Synthesis Kit (Invitrogen, Thermo Fisher Scientific). Quantitative PCR was performed with the Applied Biosystems^TM^ PowerUP™ SYBR™ Green Master Mix from Applied Biosystems on a real-time PCR instrument (LightCycler^®^ 480 System, Roche) using the primers listed in Supplementary Table 2.

### Cell transfection

Cx43 was overexpressed, as previously described^4^, in the T/C-28a2 chondrocyte cell after transfection with a pIRESpuro2 plasmid construct (Clontech) containing the human Cx43 sequence, kindly provided by Arantxa Tabernero (INCL, University of Salamanca, Spain). Electroporation was performed with the Amaxa^®^ Cell Line Nucleofector^®^ Kit V (Lonza) in a Nucleofector™ 2b device (Lonza) following the manufacturer’s instructions. Transfected cells selection was performed by culturing them with 0.1 *μ*g/ml puromycin dihydrochloride (Tocris).

### Luciferase reporter assay

A DNA construct containing the upstream 300 bp-regulatory sequence of the human Cx43 promoter (−108, +279, relative to the human Cx43 transcription start site) in a pGL3-Basic vector was kindly donated by Dr. Mustapha Kandouz (Wayne State University, School of Medicine, Detroit, USA). The T/C-28a2 chondrocyte cell line was transfected with 3 *μ*g of the plasmid using the Amaxa^®^ Cell Line Nucleofector^®^ Kit V in a Nucleofector™ 2b device. After 24 h, cells were treated with 10 *μ*M oleuropein and/or 5 *μ*g/ml of the ATP synthase blocker oligomycin (Sigma-Aldrich, O4876) for 1 h. For the luminescence analysis, the Firefly Luciferase Assay Kit from Biotium was used according to the manufacturer’s instructions. Briefly, cells were lysed with the Firefly Luciferase Lysis Buffer and the lysate was incubated with a 0.2 mg/ml luciferin solution. Firefly luminescence was measured in an Infinite 200 PRO Tecan plate reader (Tecan). Luminescence was normalized to the total protein content.

### Statistical analysis

Data were analysed using GraphPad Prism software (version 5.00). Unless otherwise indicated, analyses were performed using either the Student’s *t*-test or the Mann–Whitney *U*-test. P<0.05 was considered statistically significant.

## Acknowledgements

This work was supported in part through funding from the Spanish Foundation for Research on Bone and Mineral Metabolism (FEIOMM), grant PRECIPITA-2015-000139 from the FECYT-Ministry of Economy and Competitiveness (to M.D.M.), grant PI16/00035 from the Health Institute ‘Carlos III’ (ISCIII, Spain), the European Regional Development Fund, ‘A way of making Europe’ from the European Union (to M.D.M.) and a grant from Xunta de Galicia IN607B 2017/21 (to M.D.M.) and pre-doctoral fellowship to M.V.-E (ED481A-2015/188). We thank members of the CellCOM group for helpful technical suggestions, María Dolores Álvarez Alvariño and Jesús Loureiro for generously collecting tissue samples in the operating room after surgery, Ángel Concha and the Biobank of A Coruña, María Vázquez and Beatriz Lema for tissue processing and micromass sectioning. Moisés Blanco for helpful advice for the statistical analysis of experimental data.

## Author contributions

M.V.-E. designed and performed the experiments, analysed the data and prepared the figures. A.V.-V. assisted with cell culture and chondrocyte isolation from patients’ cartilage. C.L.P., M.K., A.C.-D. and A.C.-C. provided drugs and hMSCs from healthy donors. V.M. analysed data. E.F. and J.R.C. contributed clinical and technical advice. J.R.C. also provided cartilage tissue from patients and healthy donors. A.B. designed, performed and helped with various aspects of the flow cytometry experiments and analysis. M.D.M. conceived, directed and supervised the study. M.V.-E. and M.D.M. wrote the manuscript with input from all co-authors. All authors reviewed the manuscript.

## Abbreviations

ACAN: aggrecan;
AM: adipogenic medium;
CBX: carbenoxolone;
CD105: endoglin;
CD166: CD166 antigen (ALCAM);
CM: chondrogenic medium;
CMT: chondrocyte-to-mesenchymal transition;
Col2A1: collagen type II;
COX-2: cyclooxygenase-2;
Cx43: connexin43;
CxREs: connexin-response elements;
ECM: extracellular matrix;
EMT: epithelial-to-mesenchymal transition;
FDG: fluorescein-di-D-galactopyranoside;
GJs: gap junctions;
GJIC: gap junction intercellular communication;
hMSCs: human mesenchymal stem cells;
IL-1β: interleukin 1 beta;
IL-6: interleukin 6;
LY: Lucifer yellow;
MMP-3: matrix metalloproteinase 3;
NF-κB: nuclear factor kappa-light-chain-enhancer of activated B cells;
OA: osteoarthritis;
OACs: osteoarthritic chondrocytes;
OE: olive extract;
OM: osteogenic medium;
OSTCN: osteocalcin;
p16: cyclin-dependent kinase inhibitor 2A;
p53: cellular tumour antigen p53;
PPARγ: peroxisome proliferator-activated receptor gamma;
RT-qPCR: real-time quantitative polymerase chain reaction;
SAβG: senescence-associated β-galactosidase activity;
SASP: senescence-associated secretory phenotype;
TNFa: tumour necrosis factor alpha;
Twist-1: twist-related protein 1.

## Notes

**Conflict of interest**. The authors have declared that no conflict of interest exists.

