## SupplementaryData for "Senolytic activity of small molecular polyphenols from olive restores chondrocyte redifferentiation and cartilage regeneration in osteoarthritis"

### Supplementary Figure Legends

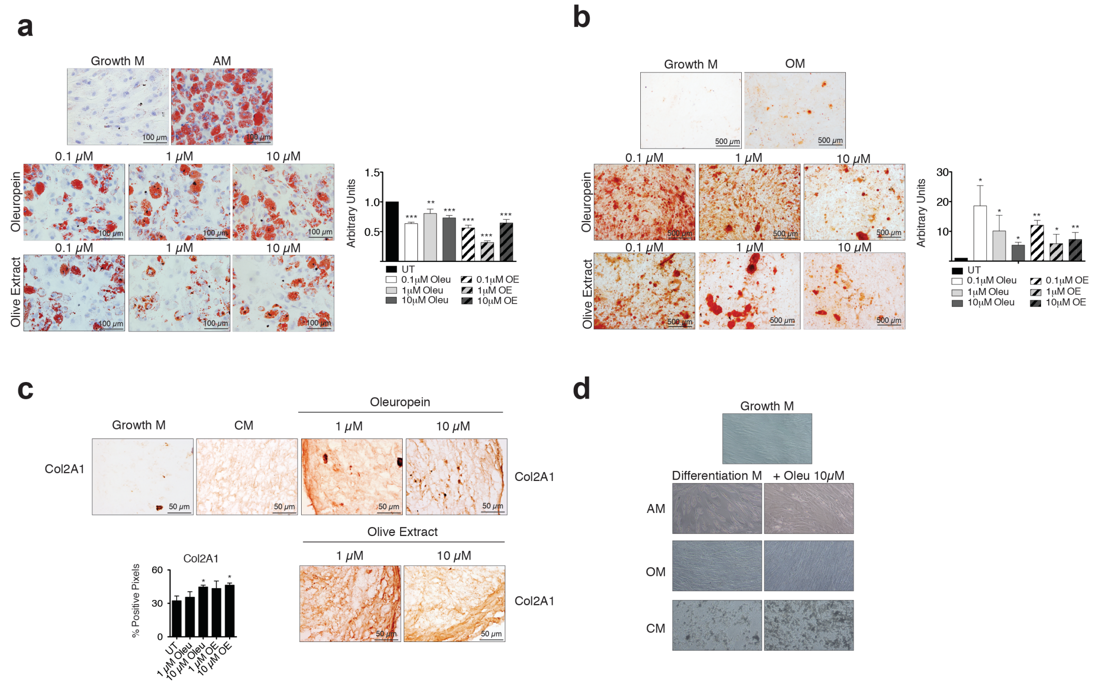


**Supplementary Figure 1.** (**a**) Adipogenic differentiation of hMSCs exposed to different concentrations of oleuropein (Oleu) or OE. Cells were stained with oil red O for lipid evaluation. The graph represents the ratio of cells with lipid deposits to the total number of cells and was normalized to hMSCs cultured in adipogenic medium (AM) (mean±s.e.m.; ***P*<0.01, ****P*<0.0001; Mann–Whitney test). (**b**) Osteogenic differentiation of hMSCs treated with Oleu or OE for 21 days. Alizarin red staining was performed to evaluate calcium deposits. Quantification was normalized to hMSCs cultured in osteogenic medium (UT) (mean±s.e.m.; **P*<0.05, ***P*<0.01; Mann–Whitney test). (**c**) Chondrogenic differentiation of hMSCs for 30 days exposed to different concentrations of Oleu or OE. Images represent Col2A1 immunohistochemistry (n=4-6; mean±s.e.m.; **P*<0.05; Mann–Whitney test). (**d**) Images showing the phenotypes of hMSCs differentiated for 14 days with or without Oleu. AM (adipogenic differentiation), OM (osteogenesis) and CM (chondrogenesis). Original magnifications ×10.

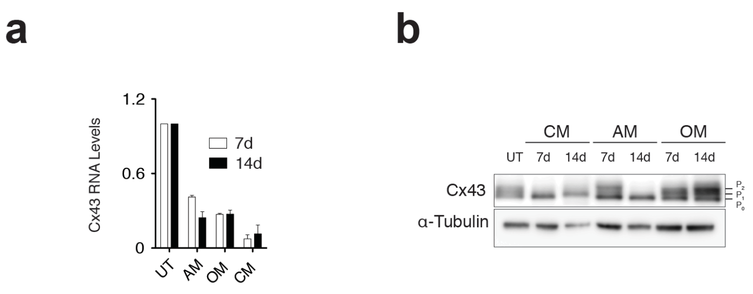


**Supplementary Figure 2.** (**a**) Cx43 levels analysed by RT-qPCR of hMSCs differentiated with adipogenic (AM), osteogenic (OM) or chondrogenic (CM) medium for 7 and 14 days (n=2-3; data were normalized to HPRT-1 levels and are shown as mean±s.e.m.). (**b**) Comparative Cx43 protein levels of hMSCs cultured in chondrogenic (CM), adipogenic (AM) and osteogenic medium (OM) for 7 and 14 days. α-tubulin was used as a loading control.

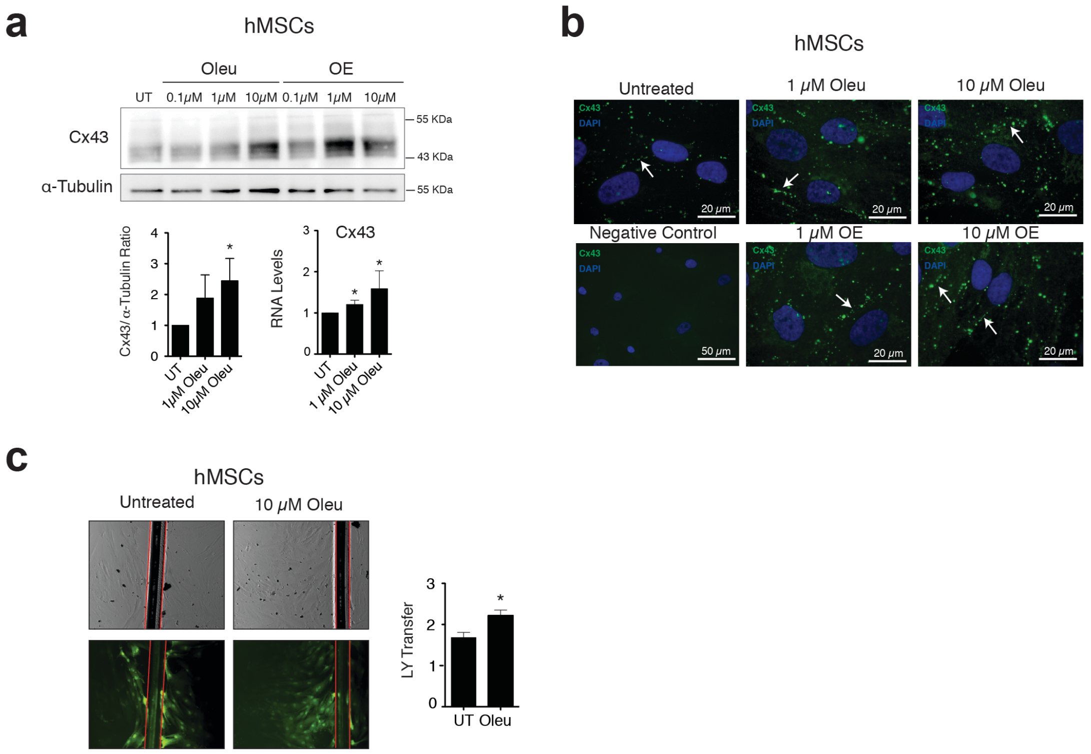


**Supplementary Figure 3.** (**a**) Western blot showing increased Cx43 protein expression in hMSCs treated for 2 h with oleuropein (Oleu) or an olive-extract (OE) containing 41.5% Oleu as well as other polyphenolic compounds. α-tubulin was used as a loading control. RT-qPCR showing increased Cx43 mRNA in hMSCs treated with Oleu in basal/growth medium (α-MEM with 10% FBS) (n=9; data were normalized to HPRT-1 levels and are shown as mean±s.e.m.; **P*<0.05; Mann–Whitney test). (**b**) Immunofluorescence analysis showing increased Cx43 at the membrane (GJ plaques, white arrows) in hMSCs treated with Oleu or OE for 2 h (growth medium). Cell nuclei were stained with DAPI. Original magnifications ×40 and ×100. Images represent two independent experiments. (**c**) SL/DT assay examining GJ activity in hMSCs treated with 10 $\mu$M Oleu or 10 $\mu$M OE. Red lines represent the cut edge where cells took up the LY immediately after scraping. The graph represents the ratio of stained cells to the number of cells at the scrape edge (LY transfer levels are represented as mean±s.e.m.; n=6 (UT) and n=15 (10 $\mu$M Oleu); **P*<0.05; Mann–Whitney test).

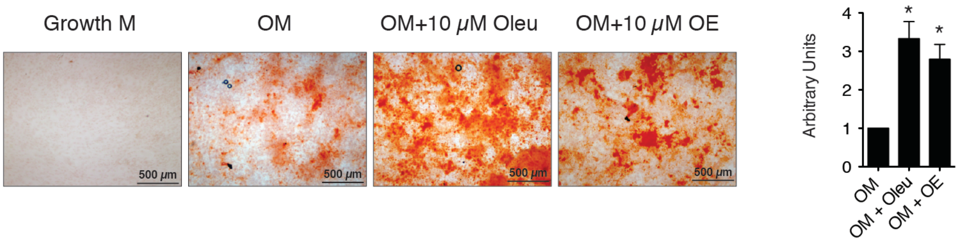


**Supplementary Figure 4.** Calcium deposits in osteogenic medium-differentiated OACs evaluated by alizarin red staining. OACs were cultured in osteogenic medium (OM) supplemented with 10 $\mu$M oleuropein (Oleu) or OE for 21 days. Quantification was performed by counting red pixels and normalization to hMSCs differentiated in osteogenic medium without treatment.

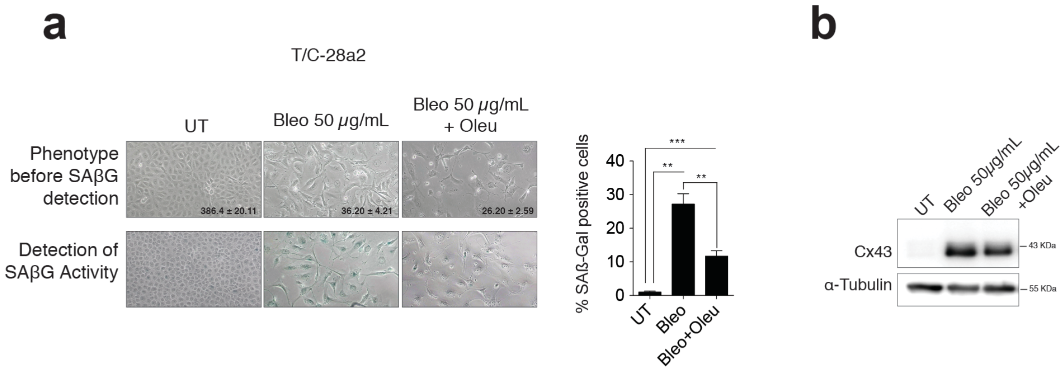


**Supplementary Figure 5.** T/C-28a2 chondrocyte cell line was cultured with bleomycin (Bleo) 50 $\mu$g/mL for 24 h, and then cultured in normal growth medium or in the presence of oleuropein 10 $\mu$g/mL for another 24 h. The mean±s.e.m of the number of cells from 5 different visual fields are shown. ß-galactosidase activity was detected by X-Gal cleavage, and cell staining (blue) was evaluated by microscopy. Quantification is shown on the right (mean±s.e.m.;** $P$<0.01, *** $P$<0.0001; Mann–Whitney test) (**a**). Cx43 levels after bleomycin or oleuropein treatments were detected by western-blot (**b**).

**Supplementary Table 1**. Listed of antibodies used in flow cytometry assays.

| Antibody | Fluorochrome | Reference | Source | Laser (nm) | Filter | Dilution |
|---|---|---|---|---|---|---|
| CD105 | PE | 105PE-100T | Immunostep | 488 | 585/42 | 1:50 |
| CD166 | APC | 1399990314 | Immunostep | 635 | 661/16 | 1:100 |
| Cx43 | APC | FAB7737A | R&D | 635 | 661/16 | 1:50 |

**Supplementary Table 2**. List of primer sequences (5'–3') used for RT-PCR analysis.

| Gene name (protein name) | Forward | Reverse |
|---|---|---|
| *ACAN* (Aggrecan) | CAGAACAACTCGGGGAACAT | GCACAATTGGAACCCTGACT |
| *Cdkn2a* ($p16^{Ink4a}$) | GAGCAGAACGATAGGGCTTG | CATGTGCCCTCTCCTCCTAA |
| *GJA1* (Cx43) | ACATGGGTGACTGGAGCGCC | ATGATCTGCAGGACCCAGAA |
| *HPRT-1* (HPRT-1) | TTGAGTTTGGAAACATCTGGAG | GCCCAAAGGGAACTGATAGTC |
| *IL-1ß* (IL-1ß) | CGAATCTCCGACCACCACTAC | TCCATGGCCACAACAACTGA |
| *IL-6* | TGTAGCCGCCCCACACA | GGATGTACCGAATTTGTTTGTA |

| | | |
|---|---|---|
| (IL-6) | | |
| *MMP-3* (MMP-3) | CCCTGGGTCTCTTTCACTCA | GCTGACAGCATCAAAGGACA |
| *CDH2* (N-cadherin) | TATTTCCATCCTGCGTGTGA | GCGTTTCATCCATACCACAA |
| *OSTCN* (Osteocalcin) | CCATGAGAGCCCTCACACTCC | GGTCAGCCAACTCGTCACAGTC |
| *PPARG* (PPARγ) | GCGATTCCTTCACTGATACACTG | GAGTGGGAGTGGTCTTCCATTAC |
| *PTGS2* (COX-2) | CTTCACGCATCAGTTTTTCAAG | TCACCGTAAATATGATTTAAGTCCA[illegible] |
| *TWIST1* (Twist-1) | CATGTCCGCGTCCCACTA | CACGCCCTGTTTCTTTGAAT |
| *VIM* (Vimentin) | ACTTTGCCGTTGAAGCTG | AATCCAGATTAGTTTCCCTCAGGT |
